# Serotonin transporter-dependent histone serotonylation in placenta contributes to the neurodevelopmental transcriptome

**DOI:** 10.1101/2023.11.14.567020

**Authors:** Jennifer C Chan, Natalia Alenina, Ashley M Cunningham, Aarthi Ramakrishnan, Li Shen, Michael Bader, Ian Maze

## Abstract

Brain development requires appropriate regulation of serotonin (5-HT) signaling from distinct tissue sources across embryogenesis. At the maternal-fetal interface, the placenta is thought to be an important contributor of offspring brain 5-HT and is critical to overall fetal health. Yet, how placental 5-HT is acquired, and the mechanisms through which 5-HT influences placental functions, are not well understood. Recently, our group identified a novel epigenetic role for 5-HT, in which 5-HT can be added to histone proteins to regulate transcription, a process called H3 serotonylation. Here, we show that H3 serotonylation undergoes dynamic regulation during placental development, corresponding to gene expression changes that are known to influence key metabolic processes. Using transgenic mice, we demonstrate that placental H3 serotonylation largely depends on 5-HT uptake by the serotonin transporter (SERT/SLC6A4). SERT deletion robustly reduces enrichment of H3 serotonylation across the placental genome, and disrupts neurodevelopmental gene networks in early embryonic brain tissues. Thus, these findings suggest a novel role for H3 serotonylation in coordinating placental transcription at the intersection of maternal physiology and offspring brain development.

## INTRODUCTION

Serotonin (5-hydroxytryptamine, 5-HT) is an essential biogenic monoamine with multipurpose functions, including regulation of fetal brain circuitry that, if disrupted, provides the foundation for behavioral dysfunction later in life^1,2^. The developing brain requires 5-HT from early embryonic stages, yet an endogenous brain-wide 5-HT source does not emerge until late in gestation^3,4^, indicating that transport of extraembryonic 5-HT to the conceptus is central to this process. Indeed, previous studies have demonstrated that the placenta, a transient endocrine and metabolic tissue at the maternal-fetal interface, delivers the majority of 5-HT into fetal circulation prior to formation of dorsal raphe nucleus projections throughout the brain^5^. Placental 5-HT may arise from different pathways, with studies describing conversion from the precursor L-tryptophan via trophoblast expression of the enzyme tryptophan hydroxylase 1 (TPH1)^6^, transporter-mediated uptake from maternal circulation via the serotonin transporter (SERT/ SLC6A4) on the placental apical membrane^7,8^, and/or regulation by the organic cation transporter 3 (OCT3/SLC22A3) at the fetoplacental endothelium^9–11^. Importantly, placental health is critical for fetal health, as indicated by numerous studies showing negative consequences on the fetal brain following placental responses to prenatal/preconception stress, inflammation, and immune activation^12–20^. Accordingly, 5-HT dysregulation also impacts vasoconstrictive properties of placental blood vessels^21,22^, as well as proliferation and viability of trophoblast cells^23^. Thus, neurodevelopment can be influenced by dysregulation of multiple 5-HT-dependent processes in placental tissues, including – but not limited to – monoamine transport. However, the mechanisms through which these 5-HT-dependent functions are regulated, as well as the modes by which placental 5-HT is acquired, are still not well understood.

Recently, a receptor-independent role for select monoamines, including 5-HT and dopamine, termed “monoaminylation,” has been described^24–27^. Monoaminylation involves the covalent attachment of free monoamine donors to glutamine-containing protein substrates by the enzyme tissue transglutaminase 2 (TGM2)^28,29^. In particular, monoaminylation using 5-HT as a donor (“serotonylation”) has been demonstrated for proteins in diverse cell types, whereby this serotonyl post-translational modification (PTM) can alter the signaling properties of bound cytosolic substrates^30–32^. In the nucleus, our group has recently demonstrated that serotonylation occurs on glutamine 5 of histone H3 (H3Q5ser)^24^. At this site, H3 serotonylation epigenetically regulates transcription either alone or in combination with the neighboring lysine 4 tri-methylation (K4me3) PTM to enhance permissive gene expression through interactions with reader proteins^33^. The combinatorial H3K4me3Q5ser PTM has been detected in regions throughout the adult brain, where it coordinates relevant gene expression programs upstream of neural differentiation and contributes to sensory processing and stress-induced behavioral plasticity in adult brain, demonstrating diverse roles for this PTM across various functional domains^34,35^. Moreover, the presence of histone serotonylation in heart, testes and other mouse organs suggest additional actions in peripheral tissues^24^. In a recent study examining human placental explants, nuclear 5-HT detected in both syncytiotrophoblasts and cytotrophoblast cells was found to be altered by inhibition of both SERT and monoamine oxidase^11^, suggesting that histone serotonylation may also be dynamically regulated in placental tissues to affect downstream processes, although follow-up studies providing evidence for this phenomenon have not yet been conducted.

Here, we investigated whether histone serotonylation may serve as an epigenetic mechanism for regulating placental gene expression programs capable of ultimately influencing offspring neurodevelopment. We found that expression of H3 serotonylation across both male and female placental development was bidirectionally regulated, with increased PTM enrichment at genomic loci related to important metabolic pathways and decreased patterns reflecting attenuation of cellular proliferation and tissue organization over development. Moreover, we demonstrate that placental 5-HT and H3 serotonylation are reliant on intact 5-HT machinery, where levels of both are reduced in tissues in which the transporters SERT, OCT3, or the enzyme TPH1 were deleted. In these tissues, we further found that SERT deletion most robustly disrupts normal H3 serotonylation patterning across the genome, with decreased enrichment at numerous loci relevant to essential placental processes. Lastly, we observed significant transcriptional abnormalities in neurodevelopmental gene networks downstream of placental changes, which appeared independent of overall 5-HT levels in brain. These findings thus establish histone serotonylation as a previously undescribed epigenetic mechanism that contributes importantly to developmental gene expression programs in placenta; phenomena that, in turn, impact key neurodevelopmental transcriptional networks in the offspring brain.

## RESULTS

### Roles for histone serotonylation in regulating gene expression programs associated with key placental functions

To begin investigating potential roles for 5-HT in placenta that could ultimately impact offspring brain development, we examined developmental 5-HT patterns occurring at E9.5 and E17.5, time points in which brain 5-HT predominantly originates from the placenta *vs.* dorsal raphe nucleus (DRN, the primary hub of 5-HTergic projection neurons in brain), respectively (**Fig. 1A**, adapted from Suri *et al.*^36^). We found that 5-HT levels in placenta decreased from E9.5 to E17.5 (**Fig. 1B**), consistent with expected 5-HT contributions from the placenta. Given our recent studies demonstrating covalent binding of 5-HT to nuclear histone proteins, we next used western blotting to assess global levels of the combinatorial serotonyl-PTM in male and female tissues at the same gestational time points. To more precisely detect fluctuations in placental 5-HT-related processes, we examined two additional time points (E12.5 and E14.5) that precede the complete formation of DRN projections throughout the embryonic brain^3,4^. We found that H3K4me3Q5ser levels decrease in placenta across gestation, with E12.5 appearing to signify the transition point after which time reductions in the mark begin to occur, with no significant effects of sex observed (**Fig. 1C, Supplementary Fig. 1**). Interestingly, the observed dynamics of histone serotonylation were also found to correspond to the extent of 5-HT supply from placenta to brain (**Fig. 1A**), suggesting that higher levels of histone serotonylation may regulate crucial placental biology at this mid-gestational window.

**Figure 1.**
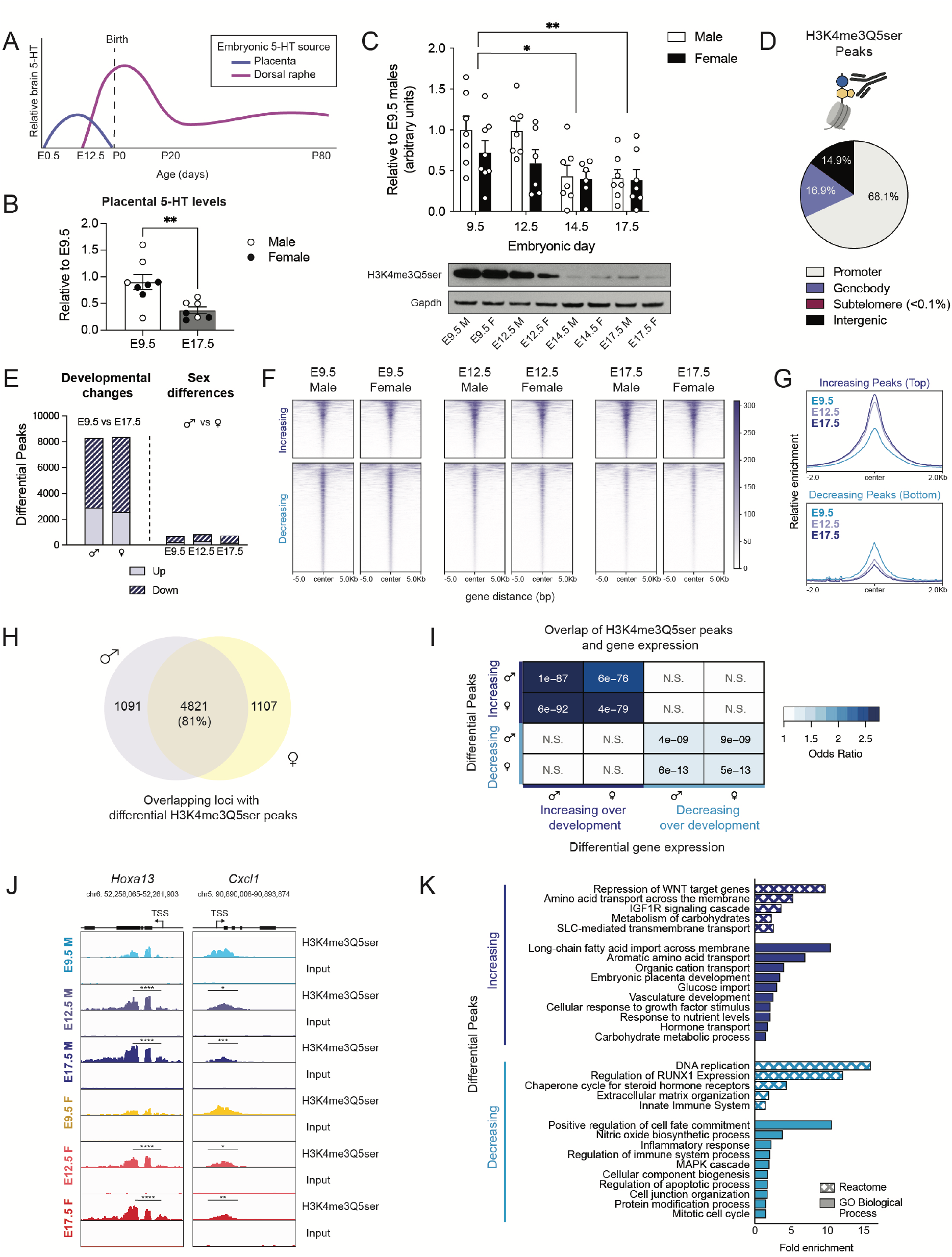
H3 serotonylation is associated with developmental gene networks in male and female placenta. **(A)** Schematic depicting brain 5-HT levels and tissue of origin, adapted from Suri *et al*.^36^ **(B)** Placental 5-HT levels decrease from E9.5 to E17.5 (unpaired Student’s t-test, t(13) = 3.209, \*\**p* = 0.0068), with male and female placental samples clustering together, as noted by circle colors (N=7-8 samples/age). **(C)** Western blot analysis of H3K4me3Q5ser in male and female placenta tissues at E9.5, E12.5, E14.5 and E17.5 showed a main effect of embryonic age (two-way ANOVA, age F(3,47) = 6.622, *p* = 0.0008) with no significant effect of sex (F(1,47) = 3.586, *p* = 0.0644), where histone serotonylation decreased over development (Sidak’s post-hoc test, E9.5 vs E14.5 (*adjusted *p* = 0.0102); E9.5 vs E17.5 (**adjusted *p* = 0.0056); E12.5 vs E14.5 (adjusted *p* = 0.057), E12.5 vs E17.5 (adjusted *p* = 0.0356), N = 6-8/group). (B, C): Data are normalized to the male E9.5 values and shown as mean ± SEM. **(D)** Averaged proportion of peaks using annotations from all developmental male and female placentas showed about 68.1% of sites found following H3K4me3Q5ser ChIP-sequencing were located in promoter regions (N = 4 samples/age/sex). **(E)** There was a ∼tenfold greater number of significantly differential peaks comparing E9.5 vs E17.5 in both males and females, compared to sex difference contrasts within embryonic age (*p* < 0.05, log_2_(fold change) > 0.1). **(F, G)** Heatmaps **(F)** and profiles **(G)** of differential peaks from E9.5 vs E17.5 comparisons, separated by directionality and centered on genomic regions to show the majority of altered peaks decrease across placental development. **(H)** Venn diagram depicting the degree of overlap between male and female E9.5 vs E17.5 comparisons using uniquely annotated peaks, indicating developmental changes are largely conserved between sex. **(I)** Odds ratio analysis of differential H3K4me3Q5ser peaks (from 1E above) and differentially expressed genes (adjusted *p* < 0.05; N = 4 samples/age/sex) from E9.5 vs E17.5 comparisons show significant association between altered histone serotonylation regulation and gene expression changes. Insert numbers indicate respective *p* values for each association (*N.S.*, *p* > 0.05). **(J)** Representative genome browser tracks of *Hoxa13* and *Cxcl1* loci for H3K4me3Q5ser (vs respective DNA input) in E9.5, E12.5 and E17.5 male and female placentas (*Hoxa13*: **** *p* < 0.0001 relative to E9.5 within sex; *Cxcl1*: \*\*\**p* < 0.001, \*\**p* < 0.01 relative to E9.5 within sex; \**p* < 0.05 denotes significant changes in E12.5 vs E17.5 males and E9.5 vs E12.5 females) Each track represents merged signal for 4 samples. **(K)** Selected Reactome and GO Biological Process pathways for differential peaks displaying significant associations with gene expression between E9.5 vs E17.5 (from 1I above) for male placenta tissues (FDR < 0.05).

As such, we next examined whether H3K4me3Q5ser is enriched at genomic loci relevant to placental functions across development. We performed chromatin immunoprecipitation followed by sequencing (ChIP-seq) in male and female placental tissues at E9.5, E12.5, and E17.5. Following peak calling in all groups, we found that the majority (∼68.1%) of H3K4me3Q5ser peaks were annotated to promoter regions, with less than a fifth of peaks each also detected in genebody and distal intergenic regions (∼16.9% and ∼14.9%, respectively; **Fig. 1D**), which is consistent with our previous findings in human neurons and rodent brain^24,35^. To identify differential enrichment sites that may regulate developmental processes, we used Diffbind to compare the earliest and latest gestational time points in our dataset^37^. In both male and female placental tissues, we identified ∼8,000 differentially enriched peaks, with the majority of these peaks for both sexes displaying significantly decreased enrichment from E9.5 to E17.5, corresponding to global western blotting patterns for the mark (**Fig. 1E, Supplementary Tables 1-2**). As the placenta is largely comprised of cells from the trophoblast lineage, which reflect fetal chromosomal sex^38^, we also examined potential sex differences in histone serotonylation. Within each developmental stage, we identified several hundred peaks altered between sexes, with E9.5 having the least (**Fig. 1E, Supplementary Tables 3-5**). Notably, at E12.5 and E17.5, the top 500 peaks showed similar sex differential patterns at the two later gestational ages, but not at E9.5, suggesting that placental sex differences in H3K4me3Q5ser enrichment are established by E12.5 and likely persist until parturition (**Supplementary Fig. 2A-C**). Annotation of these altered peaks identified sex differential sites throughout the chromosomal complement, with ∼5% located on the X and Y chromosomes (**Supplementary Fig. 2D-E**).

Given the aforementioned patterns, we next evaluated whether developmental changes in placental histone serotonylation were also impacted by sex. Hierarchical clustering of the top 1,000 peaks found to be altered between E9.5 and E17.5 revealed two sets of histone serotonylation changes (up *vs.* down), with both developmental increases and reductions from E9.5 to E17.5 displaying intermediate enrichment at E12.5, that were similarly expressed in males and females within each time point (**Supplementary Fig. 3A-B**). Visualization of all 8,274 differential histone serotonylation peaks between E9.5 *vs.* E17.5 males showed similar enrichment patterns in female placental tissues (**Fig. 1F-G, Supplementary Fig. 3C**). Comparing the degree of overlap between differential developmental sites following peak annotation, we observed an ∼81% overlap of enriched loci between males and females, altogether suggesting that these developmental changes are largely conserved between sexes in placenta (**Fig. 1H**). We next performed bulk RNA-sequencing to explore the relationship between histone serotonylation changes and gene expression in placenta. In doing so, we identified positive and significant correlations between differential gene expression and changes in serotonylation enrichment across development (**Fig. 1I, Supplementary Tables 6-8**). We observed greater transcription of gene loci with increasing H3K4me3Q5ser enrichment, as exemplified by the *Hoxa13* locus, a transcription factor critical for labyrinth vessel formation crucial for gas and nutrient exchange at the maternal-fetal interface^39^ (**Fig. 1J, Supplementary Fig. 3D**). Similarly, decreasing H3K4me3Q5ser enrichment was found to correspond to reduced gene expression, as exemplified by the *Cxcl1* locus, a chemokine ligand participant in the unique immune milieu surrounding the allogenic fetal microenvironment^40,41^ (**Fig. 1J, Supplementary Fig. 3E**). Altogether, these data indicate that H3K4me3Q5ser likely facilitates permissive transcription in placenta, similar to that of our previous findings in neural cells^24^. Functional annotation analyses (Reactome, GO Biological Process) of those loci overlapping at sites of H3K4me3Q5ser enrichment and gene expression changes (i.e., from **Fig. 1I**) further uncovered relevant gene sets to placental biology, including upregulation of vasculature development, nutrient and hormone transport processes over developmental age, and reductions in proliferative, differentiation, and immune processes near gestational term (**Fig. 1J, Supplementary Tables 9-10**)^42^.

### Placental serotonin levels are mediated by transporter-dependent pathways

Given suggestive roles of histone serotonylation in regulating the placental transcriptome, we next aimed to understand the source of its intracellular 5-HT donor pool. Prior studies have suggested several potential modes: 1) transporter-dependent mechanisms, via the high-affinity, low-capacity 5-HT uptake transporter encoded by the *Slc6a4* gene, SERT and/or the extra-neuronal organic cation transporter OCT3 (encoded by the *Slc22a3* gene), which is capable of bidirectional facultative monoamine diffusion^43,44^; or 2) intrinsic synthesis from tryptophan via trophoblast expression of TPH1^6^ (**Fig. 2A**). To assess the possibility of active 5-HT acquisition, which may serve as the donor source for the serotonyl-PTM, we chose to evaluate placental tissues at E12.5 given that H3K4me3Q5ser levels are dynamically changing between E9.5 and E17.5 to regulate placental transcriptional processes, and given the formation of a fully differentiated placenta at this stage^45^. First, to test whether placental 5-HT is transporter-mediated, we took a bioorthogonal metabolic-labelling approach, using propargylated (i.e., alkynylated) serotonin (5-PT) that allows for the immunoprecipitation of 5-PT labelled protein substrates following tissue delivery. Given prior work demonstrating that placental 5-HT depends on SERT function^7,11^, we hypothesized that 5-PT would similarly be taken up from maternal circulation via SERT. Thus, pregnant mice were injected with 100 nM or 1 μM 5-PT, based upon a reported range of 5-HT levels between basal levels *vs.* those at sites of thrombosis^46^, and conceptuses were removed 1 hour post-injection for assessments of 5-PT uptake (**Supplementary Fig. 4A**). We observed dose-dependent signals of 5-PT-labelled H3 protein in placental extracts (**Supplementary Fig. 4B**), supporting the hypothesis that histone serotonylation depends on transporter-mediated uptake of 5-HT. Subsequently, we verified placental gene expression of *Slc6a4* at E12.5 (**Fig. 2B**), also observing expression of *Slc22a3* (**Fig. 2C**), but not *Tph1* (**Fig. 2D**), further substantiating our prediction that placental 5-HT is obtained via transporters and is not endogenously synthesized. Notably, while TPH1 is not involved in placental 5-HT generation, global TPH1 knockout (KO) results in an ∼80% reduction in circulating 5-HT, which might therefore reduce the availability of 5-HT that could be taken up from circulation^47,48^.

**Figure 2.**
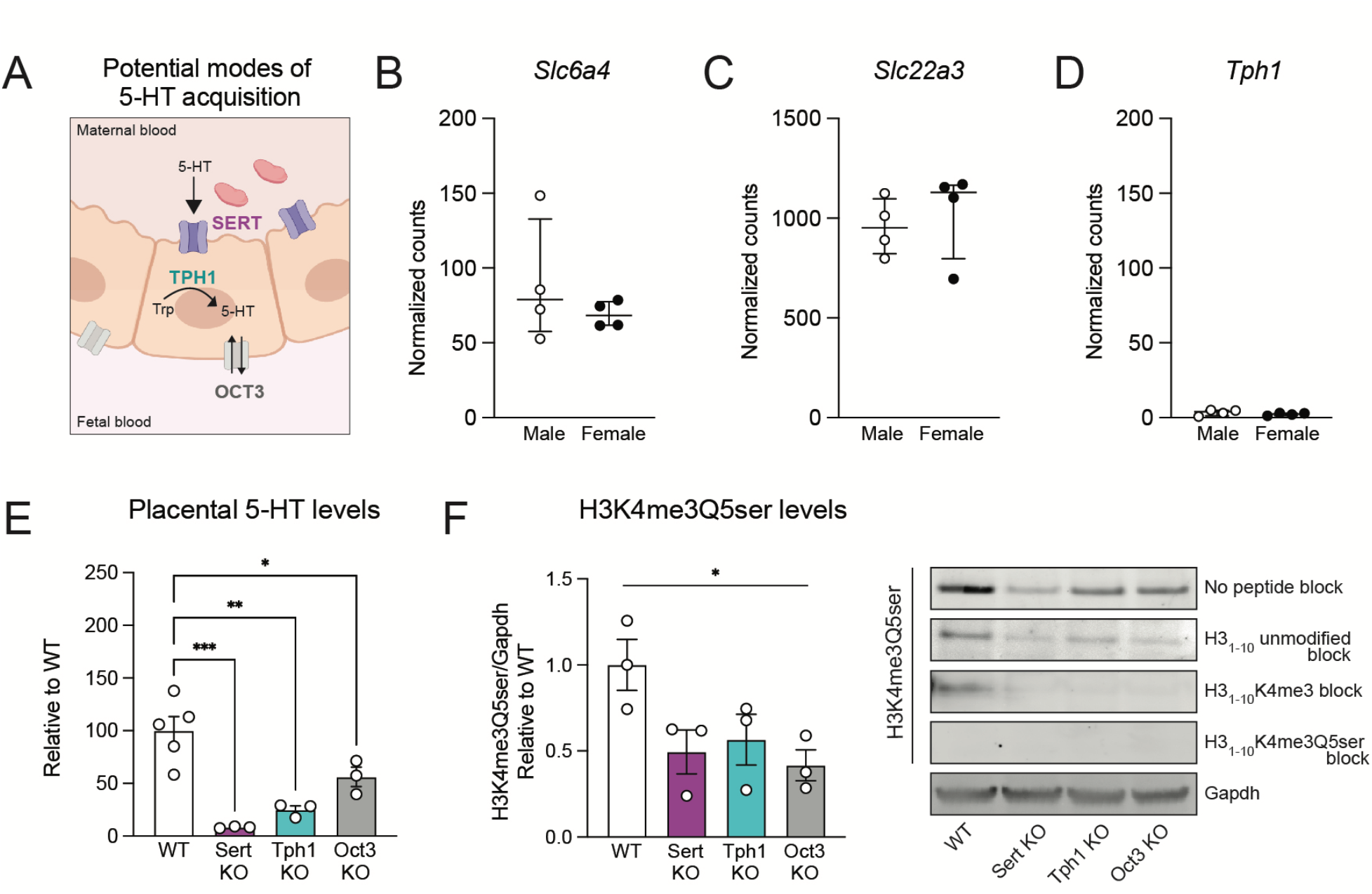
Placental 5-HT is dependent on SERT-mediated uptake. **(A)** Schematic depicting potential modes of placental 5-HT acquisition examined in this study. **(B, C)** Normalized counts indicating *Sert* (*Slc6a4*) and *Oct3* (*Slc22a3*) are expressed in both male and female placental tissues at E12.5, with no differences by sex (unpaired Student’s t-test; *Slc6a4: p* = 0.3677; *Slc22a3: p* = 0.5973). **(D)** The *Tph1* gene is not expressed in E12.5 placental tissues. N=4 samples/sex. Data are median ± interquartile range. **(E)** Assessment of 5-HT levels in E12.5 placental tissues shows significant reductions (one-way ANOVA, F(3,8) = 4.001, *p* = 0.0004) in Sert KO (Dunnett’s multiple comparisons test; ***adjusted *p* = 0.0003), Tph1 KO (**adjusted *p* = 0.0015), and Oct3 KO (*adjusted *p* = 0.04) tissues. N=3-5/group. (F) Western blot analysis of placental tissues at E12.5, showing reduced H3K4me3Q5ser in Sert KO, Tph1 KO and Oct3 KO tissues (one-way ANOVA, F(3,10) = 15.37, \**p* = 0.05). Peptide competition assays using H3_1-10_ peptides show selective signal of the serotonyl-PTM epitope is predominantly observed in WT placenta. N = 3/group. Data are mean ± SEM.

To next establish the necessity of transporters for placental 5-HT uptake and histone serotonylation deposition, we utilized transgenic mouse lines with targeted genetic deletions of *Slc6a4*, *Slc22a3* or *Tph1*. We identified robust 5-HT reductions in placental tissues from all transgenic lines examined, with the greatest loss in 5-HT signal observed in Sert KO tissues (∼90%), followed by around 70% reduction of placental 5-HT levels in Tph1 KO, and around 50% reduction in Oct3 KO (**Fig. 2E**). Thus, we next tested for corresponding reductions in global histone serotonylation levels. Indeed, western blotting revealed overall decreases in H3K4me3Q5ser signal in all three KO lines, which was further confirmed following competition assays with an H3_1-10_ peptide containing the K4me3 PTM (**Fig. 2F, Supplementary Fig. 5**). In sum, these data demonstrate placental H3 serotonylation’s reliance on 5-HT levels and the integrity of pathways regulating 5-HT entry into this tissue.

### SERT deletion downregulates histone serotonylation and disrupts developmental processes in placenta

Given histone serotonylation’s dependency on 5-HT transporter function, we next investigated whether knockout of these proteins might alter H3K4me3Q5ser enrichment at key genomic loci known to regulate placental development. Differential peak analysis following ChIP-seq demonstrated that the majority of H3K4me3Q5ser enrichment alterations observed in Sert KO, Tph1 KO, and Oct3 KO placental tissues were decreased compared to age-matched WT controls (**Fig. 3A, Supplementary Tables 11-13**). To ensure the specificity of H3K4me3Q5ser changes, we additionally performed ChIP-seq for the H3K4me3 mark alone (note that the antibody for H3K4me3 may recognize H3K4me3 both in the presence or absence of H3Q5ser^24^), which produced a distinct pattern of peak enrichment changes (**Fig. 3A, Supplementary Tables 14-16**), supporting the notion that histone serotonylation is dependent on tissue 5-HT changes rather than changes in H3K4me3 itself. Consistent with its robust 5-HT reductions, Sert KO similarly had the greatest impact on histone serotonylation peak reductions compared to deletion of OCT3 or TPH1 (**Fig. 3B, Supplementary Fig. 6**). We next evaluated the extent of overlap between developmentally relevant H3K4me3Q5ser loci that exhibit increased or decreased enrichment over embryonic age (from **Fig. 1**) with transgenic-mediated reductions in H3K4me3Q5ser or H3K4me3 enrichment. In all KO tissues, H3K4me3Q5ser-enriched loci had significantly greater overlap compared to H3K4me3 alone, with the highest degree of overlap observed for peaks altered by Sert KO (**Fig. 3C**). Therefore, we next examined those histone serotonylation peaks enriched at genomic loci at the intersection of Sert KO reductions and developmental changes occuring from E9.5 to E17.5. As expected, Sert KO downregulated H3K4me3Q5ser enrichment at these developmentally relevant loci compared to WT, Tph1 KO, and Oct3 KO placental tissues (**Fig. 3D-E**), as exemplified by the *Hoxa13* and *Cxcl1* loci (**Fig. 3F**). Functional annotation analyses of overlapping H3K4me3Q5ser-enriched loci between these multiple datasets demonstrated that SERT and TPH1 (but not OCT3) deletion disrupted important pathways for placental development, including changes in vasculature development, apoptosis, cell differentiation, and immune system processes (**Fig. 3G, Supplementary Tables 17-19**). In sum, our genomic data indicate that key moderators of the placental 5-HT donor pool lie upstream of histone serotonylation regulation. In particular, we provide evidence that SERT deletion disrupts H3K4me3Q5ser regulation of placental biology that might subsequently impact offspring brain development.

**Figure 3.**
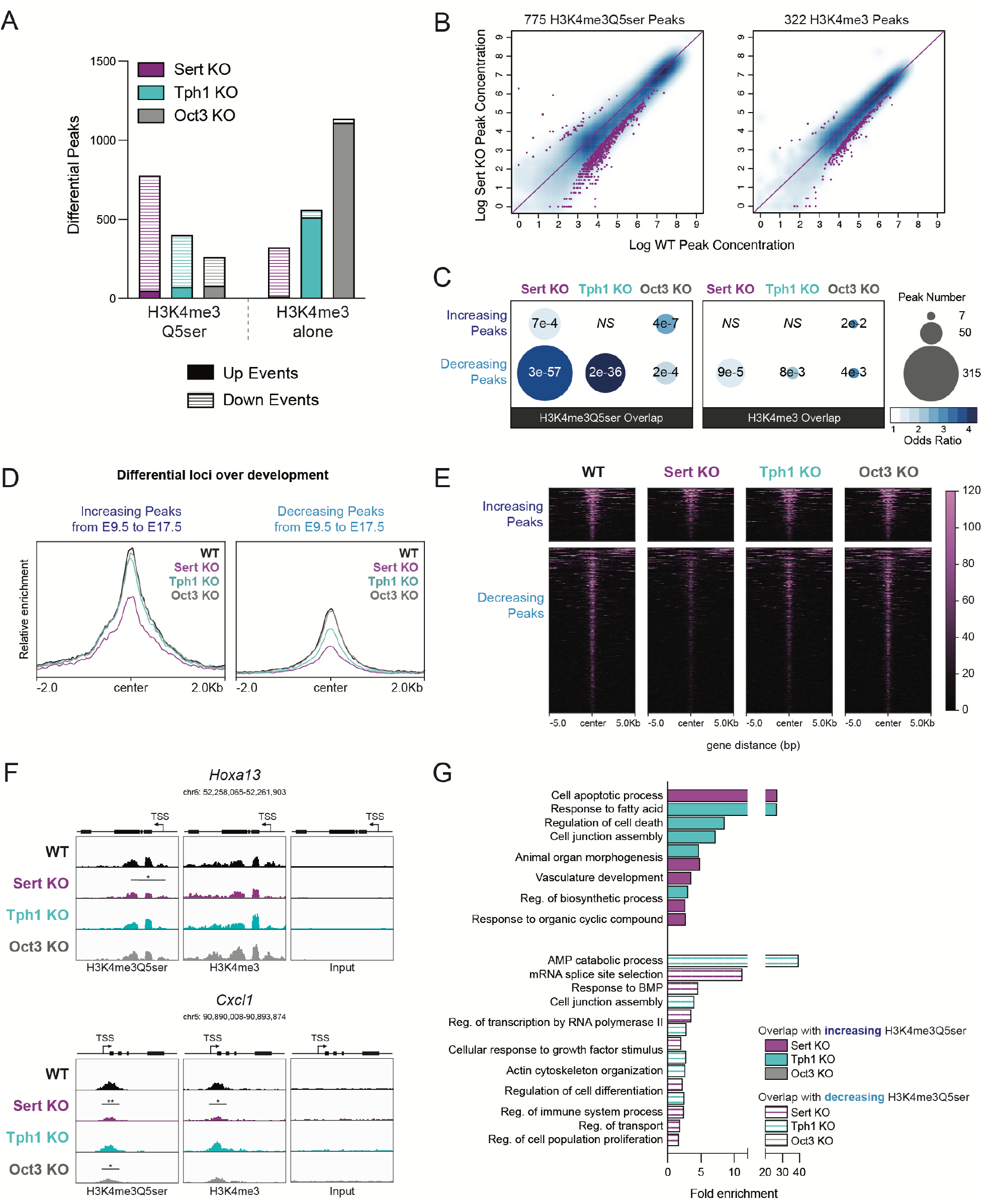
SERT deletion alters placental H3 serotonylation patterning. **(A)** Relative to WT, the greatest number of significantly decreased H3K4me3Q5ser peaks was observed in Sert KO placentas, followed by Tph1 KO and Oct3 KO (left; *p* < 0.05, log_2_(fold change) > 0.1), where the overall pattern of differential sites diverged from those of H3K4me3 alone (right; N = 3 samples/group). **(B)** Scatter plots of differential H3K4me3Q5ser (left) and H3K4me3 (right) peaks in Sert KO placentas relative to WT, showing the majority of affected sites are downregulated. **(C)** Odds ratio analysis examining overlap of significantly reduced H3K4me3Q5ser and H3K4me3 peaks (relative to WT, from 3A) with differential H3K4me3Q5ser sites between E9.5 and E17.5 (from 1E), with bubble size representing number of overlapping loci, indicating SERT deletion has greatest impact on developmentally-regulated sites. Insert numbers denote respective *p* values for each association (*NS*, *p* > 0.05), **(D, E)** Heatmaps **(D)** and profiles **(E)** of differential H3K4me3Q5ser loci between E9.5 and E17.5 that are significantly downregulated in Sert KO placentas, separated by directional changes across development and centered on genomic features. **(F)** Representative genome browser tracks of *Hoxa13* and *Cxcl1* loci for H3K4me3Q5ser and H3K4me3 (vs respective DNA input) in WT, Sert KO, Tph1 KO and Oct3 KO placentas (*Hoxa13*: \**p* < 0.05 relative to WT; *Cxcl1*: \*\**p* < 0.01, \**p* < 0.05 relative to WT for each histone modification). Each track represents merged signal for 3 samples. **(G)** Selected Reactome and GO Biological Process pathways for differential loci (vs WT) overlapping with developmentally regulated H3K4me3Q5ser sites (from 1H). Note: there were no significant pathways enriched for overlapping differential peaks from WT vs Oct3 KO comparisons (FDR < 0.05).

### Placental 5-HT and histone serotonylation reductions are associated with changes in neurodevelopmental gene expression programs

Given that the placenta is the major 5-HT source from early-to-mid gestation, we next sought to understand how brain 5-HT levels might be impacted by these placental changes (**Fig. 4A**). Importantly, the tissues used were obtained from conventional KO mice; thus we first interrogated whether transgenic-mediated changes alone might impact brain 5-HT. Transcriptomic analysis of embryonic brain tissues showed low levels of *Slc6a4, Slc22a3* and *Tph1* at E12.5 in WT mice (**Fig. 4B**), suggesting that SERT and OCT3 are not the major modes of 5-HT entry into the embryonic brain. We also examined gene expression for the neuronal isoform of tryptophan hydroxylase, TPH2 (*Tph2*), organic cation transporter OCT2 (*Slc22a2*), and the plasma membrane monoamine transporter PMAT (*Slc29a4*) to uncover other potential routes through which 5-HT in brain may be incorporated (**Fig. 4B**). Our data suggest that the E12.5 brain does not express machinery for 5-HT synthesis at this time, indicating that brain 5-HT is likely extrinsically regulated and its uptake may be mediated by the transporter PMAT, as suggested by its high levels of expression. Given that we did not observe significant expression of brain *Slc6a4, Slc22a3,* or *Tph1*, which might confound our assessments of placental 5-HT and histone serotonylation effects, we next examined how these placental disruptions might influence brain 5-HT levels. Remarkably, we observed no differences in 5-HT in any KO brain tissues compared to WT (**Fig. 4C**), similar to other studies^6,49^. We further examined whether there may be downstream differences in brain H3K4me3Q5ser abundance, but we observed no differences in any KO comparisons *vs.* WT (**Fig. 4D**, **Supplementary Figure 7**). These findings suggest that placental disruptions in 5-HT uptake do not exert direct programming effects in offspring via reductions in 5-HT delivery to the developing brain.

**Figure 4.**
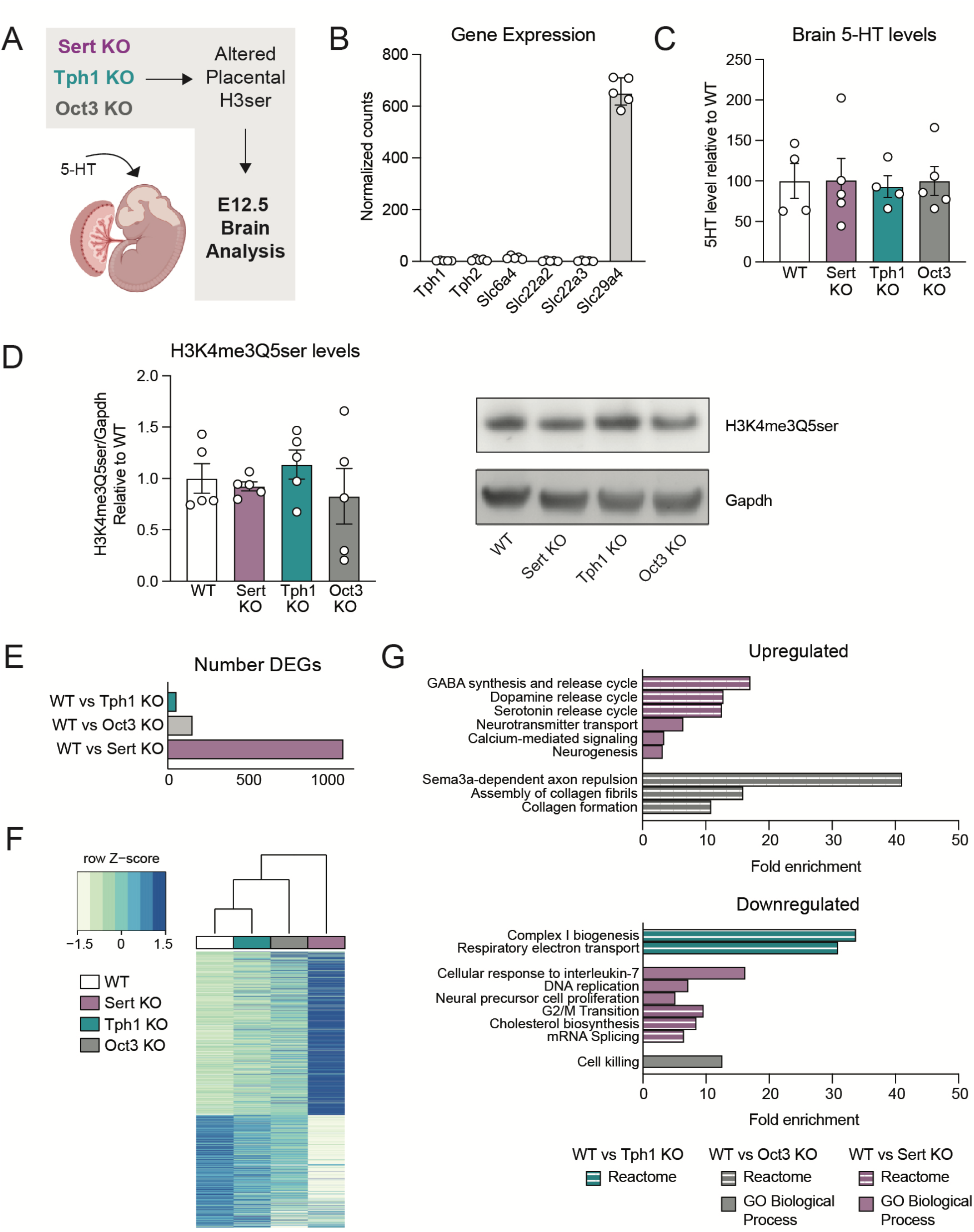
Offspring neurodevelopmental gene expression changes are associated with placental disruptions. **(A)** Schematics of study design for investigating E12.5 offspring brain changes. **(B)** Normalized counts showing gene expression for *Tph1, Slc6a4*, and *Slc22a3* are low compared to that for the transporter PMAT (*Slc29a4*) in embryonic brain. **(C)** There is no change in 5-HT levels in E12.5 brains when comparing WT vs KO tissues (one-way ANOVA, F(3,14) = 0.027, *p* = 0.9938). N=4-5 samples/group. **(D)** There also are no differences in H3K4me3Q5ser in brain tissues (one-way ANOVA, F(3,16) = 0.5861, *p* = 0.6328). N=5 samples/group. Data are mean ± SEM. **(E)** Number of differentially expressed genes from bulk RNA-sequencing comparing WT vs. Sert KO, WT vs. Tph1 KO, WT vs. Oct3 KO brain tissues at E12.5 (adjusted *p* < 0.05). **(F)** Hierarchical clustering of all differentially expressed genes relative to WT (adjusted *p* < 0.05). Expression values are averaged within genotype (N=5-6 samples/group). **(G)** Selected Reactome and GO Biological Process pathways enriched from differentially expressed genes comparing WT vs KO brain tissues at E12.5 (FDR < 0.05).

Given that SERT and TPH1 deletion both resulted in reduced placental H3K4me3Q5ser enrichment at loci involved in biosynthesis, transport, and vasculature development, we speculated that histone serotonylation might alter other placental functions that could influence the embryonic brain in a 5-HT-independent manner. Thus, to determine the overall impact of such changes on neurodevelopment, we examined embryonic brain tissues using bulk RNA-sequencing at E12.5, a time point that we already established is largely unaffected by transgenic manipulations within the brain itself. We found that the brain transcriptome was robustly altered in Sert KO tissues, with Oct3 KO and Tph1 KO brains also displaying significant regulation (all relative to WT), though to a lesser extent (**Fig. 4E-F, Supplementary Tables 20-23**). To understand what processes may be impacted in the developing brain, we performed functional annotation analyses (using GO Biological Process and Reactome databases) on differentially expressed genes from all WT *vs*. KO comparisons. Examining all significantly enriched pathways, we used Revigo to summarize redundant GO terms^50^, revealing numerous gene sets related to synaptic signaling, monoamine and neurotransmitter regulation, and neuronal proliferation altered in Sert KO brains (**Fig. 4G, Supplementary Tables 24-25**). There were also significant changes to pathways observed related to collagen formation and apoptosis in Oct3 KO brains, and downregulation of cellular respiration in Tph1 KO brains, which may be indicative of insufficient ‘fuel’ being transported from placenta to the conceptus (**Fig. 4G, Supplementary Tables 26-28)**. In total, these data indicate that even moderate changes to placental 5-HT and histone serotonylation levels appear sufficient to affect important neurodevelopmental processes in the developing fetus.

## DISCUSSION

Here, we demonstrated that histone serotonylation likely influences embryonic brain development via epigenetic regulation of the extra-embryonic placental transcriptome. We showed that H3 serotonylation is bidirectionally regulated across embryogenesis, corresponding with gene expression changes and coordination of known placental pathways that are crucial to fetal growth. We further established that SERT is the major mode of 5-HT transport from maternal peripheral circulation to placenta, a process that when disrupted also perturbs normal developmental serotonyl-PTM patterning. Moreover, we found that such disruptions in placental histone serotonylation may have important downstream effects on the embryonic brain transcriptome, supporting placental epigenetics as an exciting mechanism of neurodevelopmental programming that may affect behavioral outcomes and/or disease risk later in life. While the current study illustrates an exciting framework by which the placental 5-HT machinery intersects with chromatin mechanisms to influence offspring outcomes, there are several limitations to the current study that deserve attention. Most notably, given our use of tissues from conventional transgenic KO mice, there may be other tissue contributions involved; however, as maternal stimuli are communicated to the fetus via placental signaling, we propose that the offspring brain outcomes are directly affected by the placental changes observed in this study. Indeed, prior work suggests that increased necrosis in Sert KO and Tph1 KO placentas occurs via 5-HT receptor signaling, which is normally terminated by SERT-mediated uptake^23^. In the current study, both SERT and TPH1 deletion were found to disrupt H3K4me3Q5ser enrichment at loci involved in cell apoptotic processes, and thus may additionally regulate this phenotype via epigenetic changes. Therefore, further studies selectively targeting histone serotonylation within the placenta will be needed to fully resolve whether such 5-HT-dependent chromatin mechanisms causally contribute to placental dysregulation and/or act in parallel with disrupted receptor signaling.

Furthermore, given the essential role of developmental 5-HT on neuronal patterning, many studies have focused on identifying the mechanism through which placental 5-HT is acquired and transferred to the offspring brain. Debates regarding this source posit that placental 5-HT may derive from a maternal origin via uptake from blood, or endogenous synthesis via metabolism of the precursor L-tryptophan^6,51,52^. Using genetic targeting of these potential 5-HT sources, our findings support maternal serotonin supply as the major determinant of 5-HT and H3 serotonylation levels in placenta. Indeed, we demonstrated that *Tph1* expression is absent in placenta, similar to other studies examining human and rodent tissues^11,53,54^. For this reason, reductions in placental H3K4me3Q5ser in Tph1 KO tissues may be explained by lowered 5-HT blood levels, due to disrupted 5-HT synthesis in enterochromaffin cells^48,55^. Therefore, overlapping H3K4me3Q5ser enrichment reductions in Sert KO *vs.* Tph1 KO tissues likely occur due to a convergence of pathways dependent on 5-HT in maternal blood. In addition to reduced placental uptake via SERT deletion, Sert KO animals have low peripheral 5-HT (due to a deficiency of platelets in taking up 5-HT^56^) as observed in Tph1 KO, which result in decreased uptake into trophoblast cells, altogether indicating that placental 5-HT is of maternal origin and is not endogenously synthesized within the placenta. Indeed, genetic deletion of SERT eliminates the majority of placental 5-HT at mid-gestation. Residual H3K4me3Q5ser signal in Sert KO tissues, then, likely result from patterning at earlier time points when other modes of 5-HT acquisition may be present (e.g., other transporters and/or transient embryonic synthesis^43,44,48,57^), or technical artifacts owing to the process of polyclonal antibody generation using H3K4me3Q5ser immunogens. To control for this technical limitation, we additionally performed H3K4me3 ChIP-sequencing and observed that while there were indeed differential sites of overlap between H3K4me3 and H3K4me3Q5ser, differential histone serotonylation could not be accounted for by changes in H3K4me3 alone. Instead, we observed that reduced H3K4me3Q5ser patterns in KO placentas closely corresponded with the extent of 5-HT decreases, suggesting that this PTM depends on donor availability (consistent with our previous biochemical analyses^58^). It is also worth noting that the overlapping reductions in signal observed between H3K4me3Q5ser and H3K4me3 alone may occur due to previous observations that H3Q5ser inhibits H3K4 demethylase activity, and thus loss of the serotonyl-PTM may additionally destabilize the presence of H3K4me3 at certain loci^59^.

The developing brain is highly sensitive to placental insults resulting from environmental perturbations and imbalances of specific nutrients, hormones, and other chemical signals^38^. Using transgenic KO mice, we identified a specific time point in which there was minimal expression of key 5-HT machinery within the brain, allowing us to examine non-cell autonomous effects originating from deletion of SERT, TPH1 or OCT3 in the placenta and/or maternal tissues. Indeed, we detected robust differential gene expression in the E12.5 Sert KO brain, supporting functional responsivity to placental effects. As previously mentioned, we must cautiously interpret these findings given the use of whole-body KO animals. Beginning at E10.5, SERT is detected in embryonic cardiac and liver tissues^60^, and it is possible that disruptions to these systems may result in excess 5-HT in fetal circulation that also contribute to brain changes. In this way, the effects observed in Tph1 KO brains, though more subtle, provide clearer proof-of-concept evidence that placental 5-HT and histone serotonylation directly impact brain programming, due to restricted non-neuronal *Tph1* expression that is not detected until E14.5^55^.

With respect to how precisely placental histone serotonylation changes may mediate brain reprogramming, we did not expect that 5-HT levels would be unaffected in the corresponding KO brains given the robust 5-HT reductions observed in Sert KO and Tph1 KO placentas, though it is notable that other studies have made similar observations^6,49^. There are several potential explanations: it is possible that the placenta buffers against 5-HT deficiencies, such that the embryo nonetheless attains the necessary amount, or there may be alternate 5-HT sources that compensate for placental insufficiency^48^. The answer to this question is beyond the scope of the current study, but will be crucial to understanding the complex role of placental 5-HT signaling in developmental brain programming. While we do not detect global histone serotonylation changes within the brain itself, this is likely due to the specific time point examined. For example, SERT expression increases across gestation and is transiently upregulated in the thalamus and hippocampus during early postnatal development, where it is critically necessary for neuronal projection patterning^61,62^. Moreover, SERT inhibition during early postnatal windows, but not in adulthood, results in behavioral deficits later in life^63^. Indeed, we postulate that histone serotonylation governs transcriptomic patterns during these select neurodevelopmental windows (as we have described previously in culture systems using neuronal precursor cells and human induced pluripotent stem cell-derived 5-HTergic neurons^24^), which are the subject of future investigations, but that during early-to-mid embryogenesis, downstream consequences of placental 5-HT disruptions are mediated by non-serotonergic processes in the brain.

Together, our findings establish that placental H3K4me3Q5ser lies at the intersection of maternal 5-HT detection, regulation of tissue transcriptional networks, and offspring brain development, though additional studies will be needed to fully delineate the specific involvement of this histone PTM in modulating tissue-specific functions. Given that the endocrine placenta dynamically regulates H3K4me3Q5ser in response to both SERT disruptions and 5-HT changes in the maternal milieu, outstanding questions regarding the effects of prenatal stress and antidepressant exposures remain. Notably, several studies examining the effects of maternal perturbations observed dysregulation of placental 5-HT^15,64–66^; therefore, understanding how these triggers may enact negative long-term outcomes on fetal development via placental histone serotonylation changes, how fetal sex impacts these outcomes, and how antidepressant usage may reverse such dysregulated processes, are needed. Moreover, while we show that H3 serotonylation is a dynamic mechanism of developmental regulation within the placenta, a comprehensive catalogue of monoaminylated proteins (including serotonylation of both nuclear and cytoplasmic substrates) and their downstream effects on offspring neurodevelopment may provide further insight into how non-canonical monoamine mechanisms contribute to origins of neurodevelopmental disease risk.

## MATERIAL AND METHODS

### Animals

Wild-type C57BL6/J mice were purchased from Jackson Laboratories at 8 weeks old, and maintained on a 12-h/12-h light/dark cycle throughout the entirety of the experiment. Mice were provided with *ad libitum* access to water and food throughout the entirety of the experiment. All animal procedures were done in accordance with NIH guidelines and with approval with the Institutional Animal Care and Use Committee of the Icahn School of Medicine at Mount Sinai. For transgenic tissue studies, wild-type (WT), TPH1-deficient (Tph1-KO)^67^, SERT-deficient (Sert-KO)^68^ (Jackson Laboratories, stock #008355) and OCT3-deficient (Oct3-KO)^69^ (provided by Dr. Ciarimboli), all on C57Bl6/N genetic background, were bred at the MDC animal facility (Berlin, Germany) in individually ventilated cages (Tecniplast, Italy) under specific pathogen-free, standardized conditions in accordance with the German Animal Protection Law. Mice were group-housed at a constant temperature of 21 ± 2°C with a humidity of 65 ± 5%, an artificial 12 hours light/dark cycle, and with free access to water *ad libitum*. All experimental procedures were performed according to the national and institutional guidelines and have been approved by responsible governmental authorities (Landesamt für Gesundheit und Soziales (*LaGeSo)*, Berlin, Germany).

### Timed Breedings

Adult virgin female mice were bred in-house with age-matched males. Copulation plugs were checked every morning within 1 hour after lights on, where confirmation of a plug was designated as E0.5 and signaled the immediate removal of the female to her own cage with a nestlet.

### Tissue Collection and Sex Determination

Timed pregnant dams were deeply anesthetized with isoflurane at designated embryonic time points, and conceptuses were isolated from the uterine wall, as previously described^65^. Placental tissues were hemisected in the transverse plane with removal of decidua cells^70^, flash frozen on dry ice, and stored at -80°C until further processing. Enriched fetal brain tissues were separated from the head by a single cut above the eye, perpendicular to the anterior-posterior axis. All tissues were flash frozen on dry ice and stored at -80°C until further analyses. Embryonic tails for WT developmental studies were retained for sex determination by *Jarid1* genotyping, as previously described^71^. For KO studies, both male and female tissues were used per genotype after determining there were no sex differences in *Slc6a4, Slc22a3,* and *Tph1* gene expression (**Fig. 2B**) and due to limited sample *n* per group.

### 5-PT Injection and Detection

5-PT was diluted in 1x PBS to 100 nM or 1 µM, representing endogenous levels of 5-HT at basal or inflammatory conditions^46^. Pregnant mice (E12.5) were injected via tail vein with 5-PT mixtures or vehicle. 1 hour post-injection, conceptuses were removed and placental tissues were collected for further processing. Magnetic streptavidin beads (Thermo Fisher 11205D) were incubated with 10 mM biotin azide (probe condition; Click Chemistry Tools 1265) or 10 mM desthio-biotin (no probe condition; Sigma D1411) on a rotator for 1 hour at 4°C. For copper-click chemistry, placental whole cell lysates containing proteins labelled with the alkyne-functionalized 5-PT were incubated with conjugated beads, 800 µM CuSO_4_, and 400 µM sodium ascorbate added in that order on a rotator for 1 hour at 4°C in a total volume of 500 µl in 1x PBS. Reactions were stopped by adding EDTA to a final concentration of 20 mM. All samples were washed on a magnetic stand using 0.1M glycine and High Salt Buffer (500mM KCl, 20 mM HEPES, 10 mM MgCl_2_, 1% NP-40). After the last wash, sample buffer was added to beads and boiled at 95°C for 10 min, followed by gel electrophoresis and incubation with appropriate primary and secondary antibodies.

### Serotonin ELISA

Placental or fetal brain tissues were homogenized in cold PBS with 1x protease inhibitor cocktail (Roche). 60 ug of lysate per sample was quantitated using the BCA Protein Assay Kit (Pierce) and mixed 1:1 with assay buffer for measurement. Tissue 5-HT levels were assessed using the Serotonin ELISA Kit according to manufacturer’s instruction (Abcam ab133053).

### Western Blotting and Antibodies

Placental or fetal brain tissues were homogenized and sonicated in cold RIPA buffer (50 mM Tris-HCl, 150 mM NaCl, 0.1% SDS, 1% NP-40) supplemented with 1x protease inhibitor cocktail (Roche). 30 ug of protein per sample was quantitated using the BCA Protein Assay Kit (Pierce) and loaded onto 4-12% NuPage BisTris gels for electrophoresis. Fast transfers were performed using the Trans-Blot Turbo Transfer System (Bio-Rad) for 7 minutes onto nitrocellulose membranes, and blocked in 5% milk or bovine serum albumin (BSA) diluted in 0.1% PBS-T. Membranes were incubated overnight with primary antibodies at 4°C on an orbital shaker. The following day, blots were washed 3x with PBS-T at room temperature, incubated for 1 hour with secondary antibody, and washed again with PBS-T 3x. Bands were detected using either enhanced chemiluminescence (ECL; Millipore) or fluorescence with the ChemiDoc Imaging System (Bio-Rad). Densitometry was used to quantify protein bands via Image J Software and proteins were normalized to total Gapdh. For developmental H3K4me3Q5ser western blots, one sample (run 2x) was removed due to lack of signal, as indicated in Supplementary Figure 1. For peptide competition assays, antibodies were pre-incubated with indicated peptides at 1:3 concentration of peptide to antibody for 1 hour at room temperature. Following pre-incubation, membranes were incubated with the designated antibody/peptide mixture overnight at 4°C on an orbital shaker. The following combinations of antibodies/buffers were used.

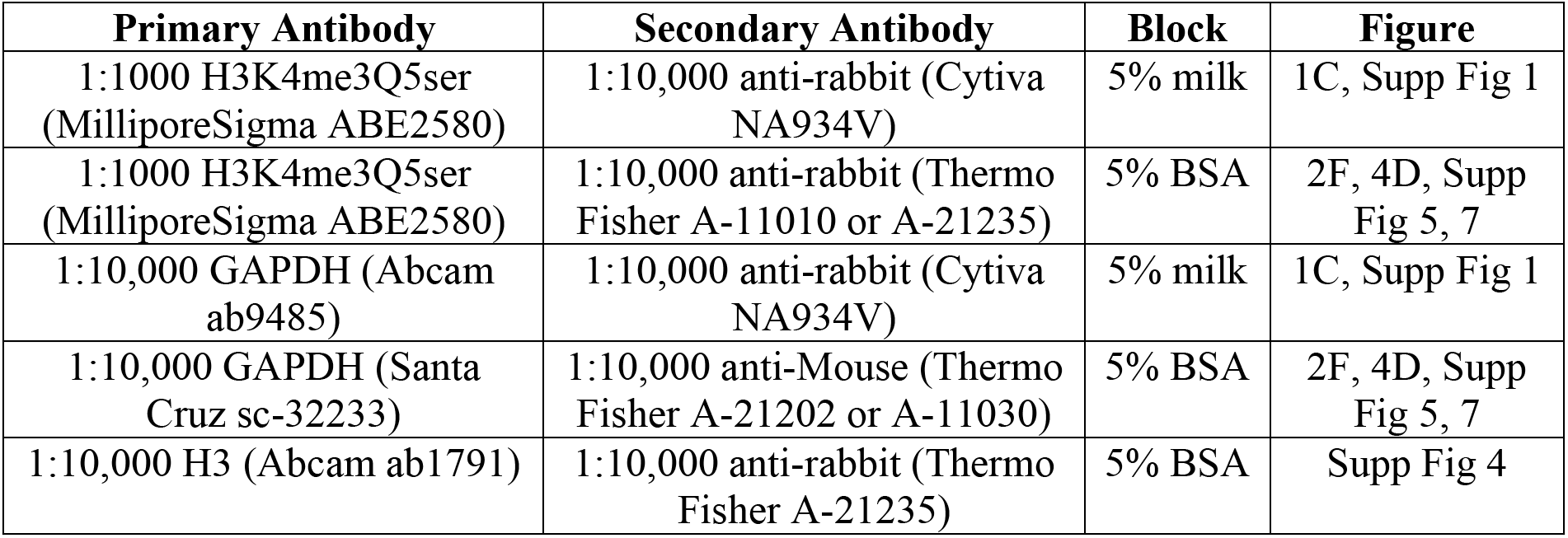

### Chromatin Immunoprecipitation, ChIP-seq and Analysis

Chromatin from hemisected placental tissues were fixed with 1% formaldehyde rotated for 12 minutes at room temperature and was subsequently quenched using a final concentration of 125mM glycine. Samples were thoroughly homogenized and washed with ice cold PBS. Fixed chromatin was sonicated using a Covaris E220 for 30-60 minutes at 4°C with the following conditions: peak incident power, 140; duty factor, 10%; Cycles/burst, 200; Water level, 0. Equal amounts of chromatin per sample were rotated with select antibodies (2.5 μg antibody/sample of either H3K4me3Q5ser (MilliporeSigma ABE2580) or H3K4me3 (Active Motif 39159)) bound to M-280 Dynabeads at 4°C overnight. The next morning, samples were washed, eluted, and reverse-crosslinked at 65°C. Samples underwent RNA and protein digestion, and DNA was purified using QIAQuick MinElute Spin columns (Qiagen 28140). 1% inputs were removed prior to antibody incubation and purified in parallel with corresponding immunoprecipitates. ChIP-seq libraries were generated using the TruSeq ChIP Library Preparation Kit (Illumina IP-202-1012) according to manufacturer’s protocol and sequenced on an Illumina HiSeq2500 or NovaSeq6000. Raw peaks were aligned to the mm10 mouse genome using the NGS Data Charmer pipeline with default settings (HISAT v.0.1.6b)^72^. Peak calling was performed using macs2 (v.2.1.1) on individual files with default settings and filtered for peaks with FDR < 0.05^73^. Differential peak analysis was conducted via pairwise comparisons using the DiffBind package (v3.8.4)^37^. Differential peaks were filtered first by log_2_(fold change) > 0.1 and defined by *p* < 0.05, where log_2_(fold change) was calculated as log2(E17.5 conc) - log2(E9.5 conc) for developmental comparisons; log2(female conc) - log2(male conc) for sex differences; and log2(KO conc) - log2(WT conc) for transgenic comparisons. These criteria were determined by visual confirmation of differential peaks after inspection of more than 100 sites in the Integrative Genomics Viewer (Broad Institute, v2.11.1). All peaks were annotated to the mm10 genome using the Homer package (v4.10)^74^. Functional annotation analysis of uniquely annotated loci was conducted using ShinyGO v0.77 with a background of all protein-coding genes in the mm10 genome^75^, with significant pathways defined by FDR < 0.05 and GO term redundancy reduction using Revigo v1.8.1^50^. Visualization of differential peaks were accomplished using internal functions of the DiffBind package or deepTools v3.5.3^76^.

### RNA Isolation, RNA-seq and Analysis

Total mRNA from hemisected placental tissues and embryonic brain tissues were extracted following homogenization in Trizol Reagent (Thermo Fisher) with subsequent clean-up using RNeasy Microcolumns (Qiagen) according to manufacturer’s recommendation. 200ng mRNA per sample was used for RNA-seq library preparation using the TruSeq RNA Library Prep Kit v2 (Illumina RS-122-2001) according to manufacturer’s protocol. Quality control of all libraries were conducted using a Qubit Fluorometer 2.0 (Thermo Fisher) and Bioanalyzer High Sensitivity DNA Analysis (Agilent) prior to sequencing on either an Illumina HiSeq2500 or NovaSeq6000. Raw fastq files containing an average of 20-30 million reads were processed for pseudoalignment and abundance quantification using Kallisto (v.0.46.1) against the EnsemblDB mus musculus (v79)^77^. To account for unwanted technical variation between batches of animal orders, sample collection, mRNA extraction, and library preparation that are each represented per sample batch, RUVs (v1.32.0) was used with a negative control gene set of total genes identified per sequencing experiment following confirmation that unwanted factors did not correlate with covariates of interest (for all experiments, k=4 was used) as previously described^78,79^. Next, differential expression analysis was performed using DESeq2 (v1.38.3) and significant genes were defined by adjusted *p* < 0.05^80^. Odds ratio overlap analysis was conducted using the GeneOverlap package (v.1.36.0), with significance indicated by *p* < 0.05. Functional annotation analysis of differentially expressed genes was performed using ShinyGO v0.77 with a background of all protein-coding genes in the mm10 genome, with significant pathways defined by FDR < 0.05 and GO term redundancy reduction using Revigo v1.8.1^50,75^. Importantly, increased *Slc6a4* expression was observed in RNA-seq data from Sert KO embryo brains, likely reflecting the aberrant introduction of an internal promoter in the design of this transgenic line and/or increased expression of transcripts that undergo nonsense-mediated decay, as indicated by loss of functional protein (Supplementary Fig. 4). Thus, to ensure nonfunctional increases in *Slc6a4* expression did not misleadingly contribute to pathway enrichment data, *Slc6a4* was removed from significant differential gene expression lists in WT vs. Sert KO comparisons prior to pathway analysis.

### Data and Materials Availability

The RNA-seq and ChIP-seq data generated in this study have been deposited in the National Center for Biotechnology Information Gene Expression Omnibus (GEO) database under accession number GSE246540. We declare that the data supporting findings for this study are available within the article and Supplementary Information. Related data are available from the corresponding author upon reasonable request. No restrictions on data availability apply.

## Supplementary Figure Captions

**Supplementary Figure 1.**
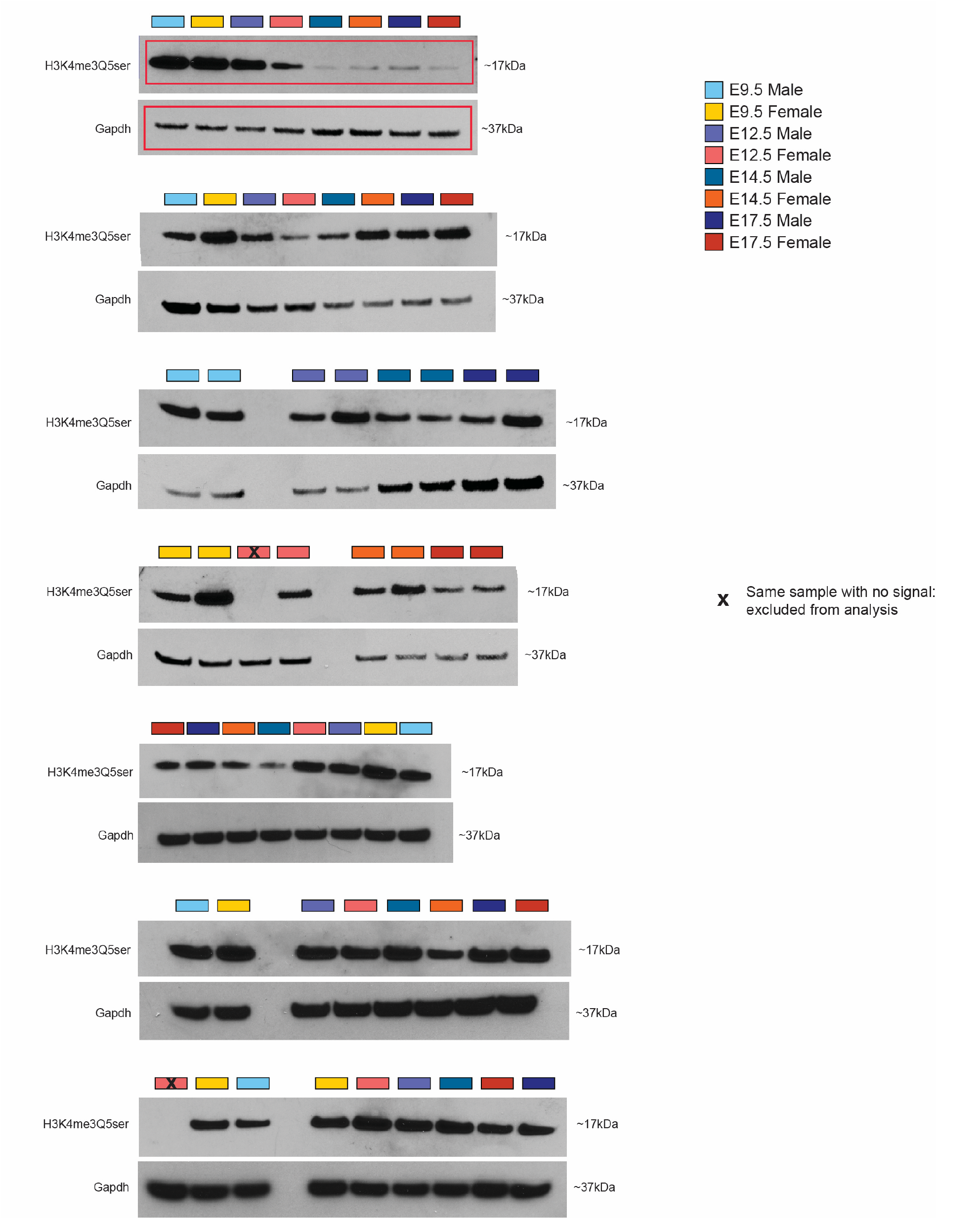
Western blots used for the quantification of the placental H3K4me3Q5ser signal in Figure 1C. Red rectangles indicate the representative blots displayed in the main figure. One sample (run two times) was excluded due to lack of H3K4me3Q5ser signal (indicated by X).

**Supplementary Figure 2.**
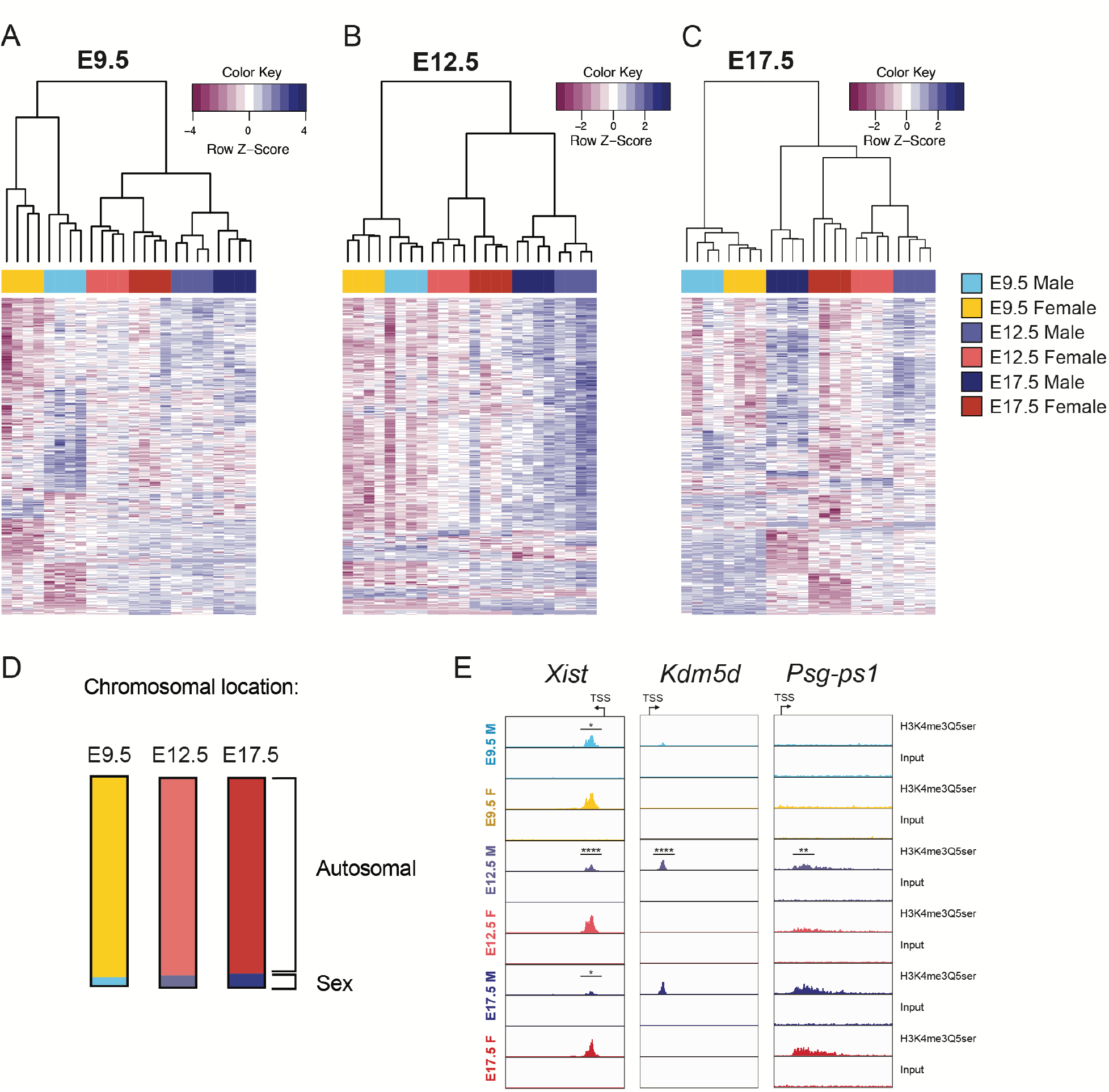
Sex differences in placental H3K4me3Q5ser (A-C) Heatmaps of top 500 differential peaks from Diffbind analysis (*p* < 0.05) based on fold change comparing male vs. female placental tissues within age at **(A)** E9.5, **(B)** E12.5, and **(C)** E17.5 with hierarchical clustering (N=4 samples/sex/age). **(D)** Chromosomal location of all significant (*p* < 0.05) sex different peaks per age, where ∼95% of peaks occurred on autosomal chromosomes and 4-6% of peaks occurred on the X or Y chromosomes. **(E)** Representative genome browser tracks of sex different H3K4me3Q5ser peaks (vs respective DNA input on the X chromosome (*Xist*: *p* < 0.05 relative to male at E9.5 and E17.5, *p* < 0.0001 relative to male at E12.5), Y chromosome (*Kdm5d*: *p* < 0.0001 relative to male at E12.5), and on chromosome 7 (*Psg- ps1*: *p* < 0.01 relative to male at E12.5). Each track represents merged signal for 4 samples.

**Supplementary Figure 3.**
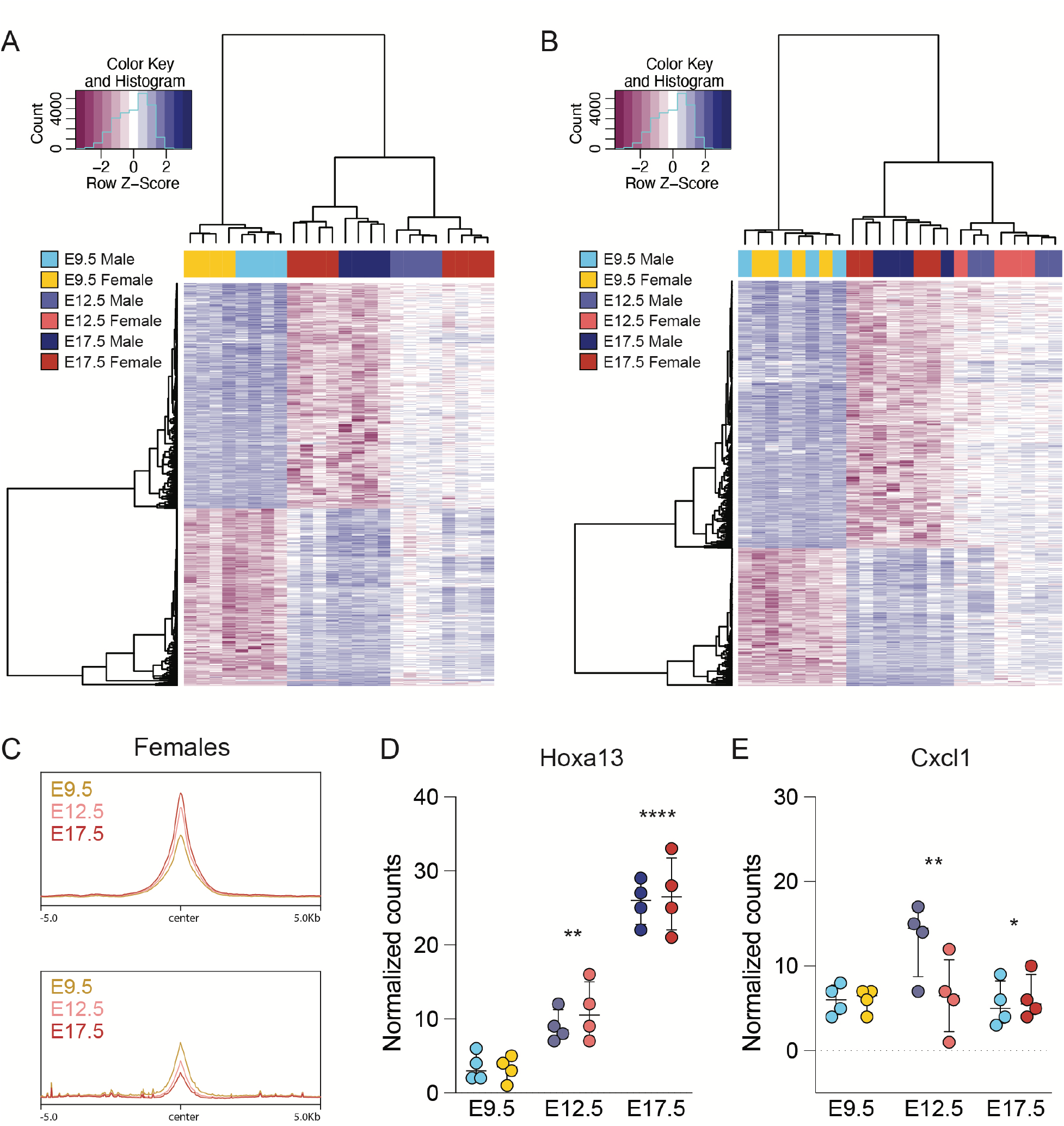
Differential H3K4me3Q5ser enrichment in male and female placenta (A-B) Heatmaps of top 1000 differential peaks from Diffbind analysis (*p* < 0.05) based on fold change comparing E9.5 vs E17.5 in male **(A)** and female **(B)** placental tissues with hierarchical clustering, showing developmental changes are largely conserved across sexes (N=4 samples/sex/age). **(C)** Profiles of differential peaks from E9.5 vs E17.5 comparisons, separated by directionality and centered on genomic regions in females. **(D)** *Hoxa13* gene expression increases across development, corresponding with H3K4me3Q5ser increases (two-way ANOVA, effect of age: F(2,18) = 1108, *p* < 0.0001; Tukey’s post-hoc, E9.5 vs E12.5: \*\**p* = 0.0016; E12.5 vs E17.5: \*\*\*\**p* < 0.0001). **(E)** *Cxcl1* gene expression decreases from E12.5 to E17.5, similarly to H3K4me3Q5ser decreases at the same time points (two-way ANOVA, effect of age: F(2,18) = 4.244, *p* = 0.0309; Tukey’s post-hoc; E12.5 vs E17.5: \**p* = 0.049). N = 4/sex/age. Data are median ± interquartile range.

**Supplementary Figure 4.**
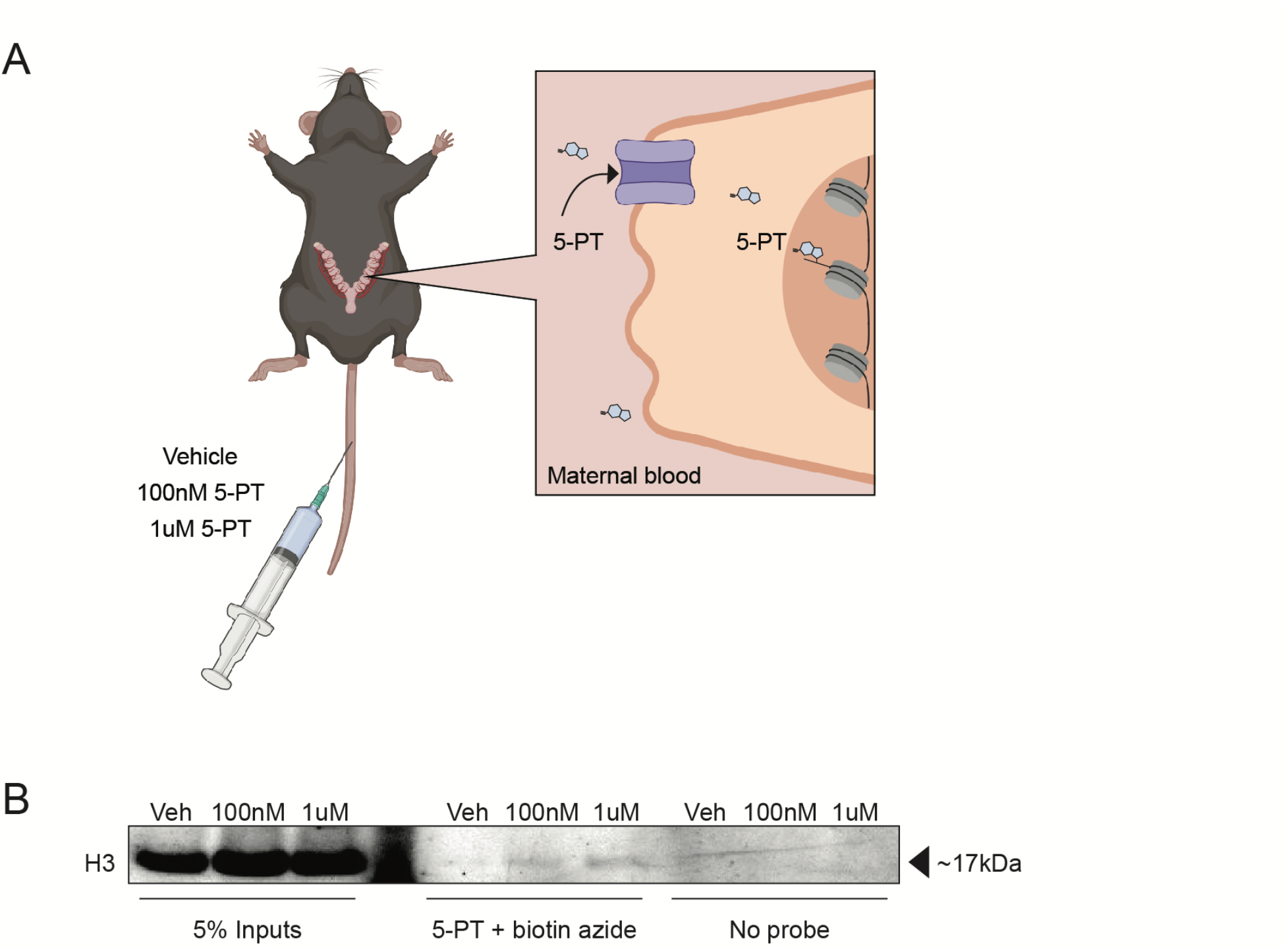
Detection of circulating propargylated 5-HT (5-PT) on placental histone H3. **(A)** Schematics of experimental design and expected results. The alkyne-functionalized 5-HT analogue, 5-PT, was injected into pregnant E12.5 mice at 100nM or 1µM (vs. vehicle). Transporter-mediated mechanisms within the apical membrane of the placenta facing maternal circulation would directly take up 5-PT into cells, increasing 5-PT addition to histone H3 as proxy for how H3 serotonylation might be regulated. One hour post-injection, placental tissues were collected and subjected to copper-click chemistry using biotin azide (vs. no probe). 5-PT-ylated proteins were immunoprecipitated with streptavidin beads, followed by western blotting for H3. **(B)** Western blot showing H3 is enriched in 5-PT-treated samples in dose-dependent manner (compared to vehicle) only in click reaction conditions.

**Supplementary Figure 5.**
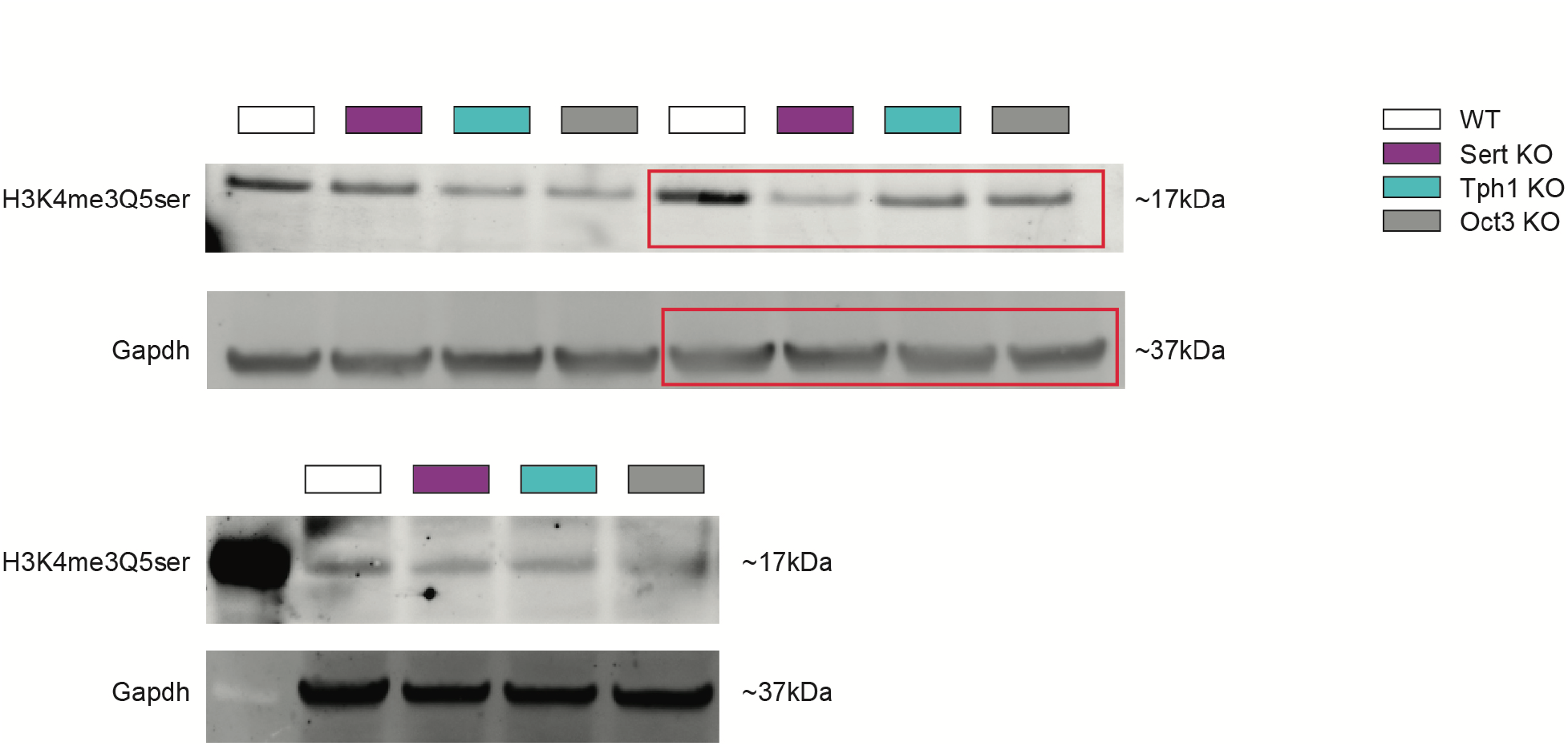
Western blots used for the quantification in Figure 2F. Western blots of placental tissues at E12.5, showing H3K4me3Q5ser in WT, Sert KO, Tph1 KO and Oct3 KO tissues. Red rectangles indicate the representative blots displayed in the main figure.

**Supplementary Figure 6.**
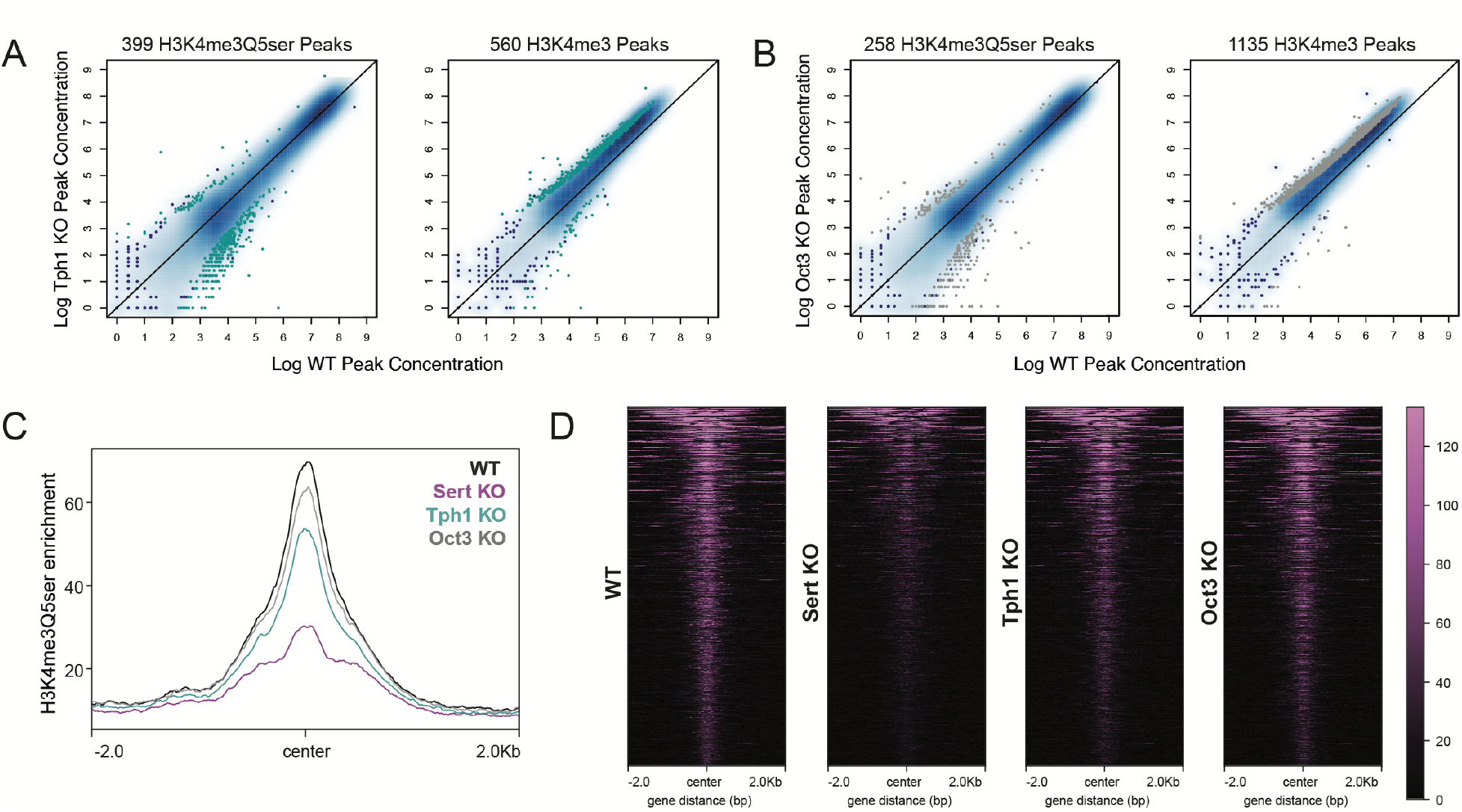
Histone serotonylation reductions are most affected by deletion of SERT (A-B) Scatterplots of differential H3K4me3Q5ser and H3K4me3 peaks in **(A)** Tph1 KO and **(B)** Oct3 KO E12.5 placentas relative to WT (*p* < 0.05). **(C)** Profiles and **(D)** heatmaps of all downregulated differential H3K4me3Q5ser loci comparing WT vs Sert KO placental tissues (*p* < 0.05), showing Sert KO has the greatest impact on histone serotonylation peak reductions compared to Tph1 KO and Oct3 KO, consistent with reductions in 5-HT levels.

**Supplementary Figure 7.**
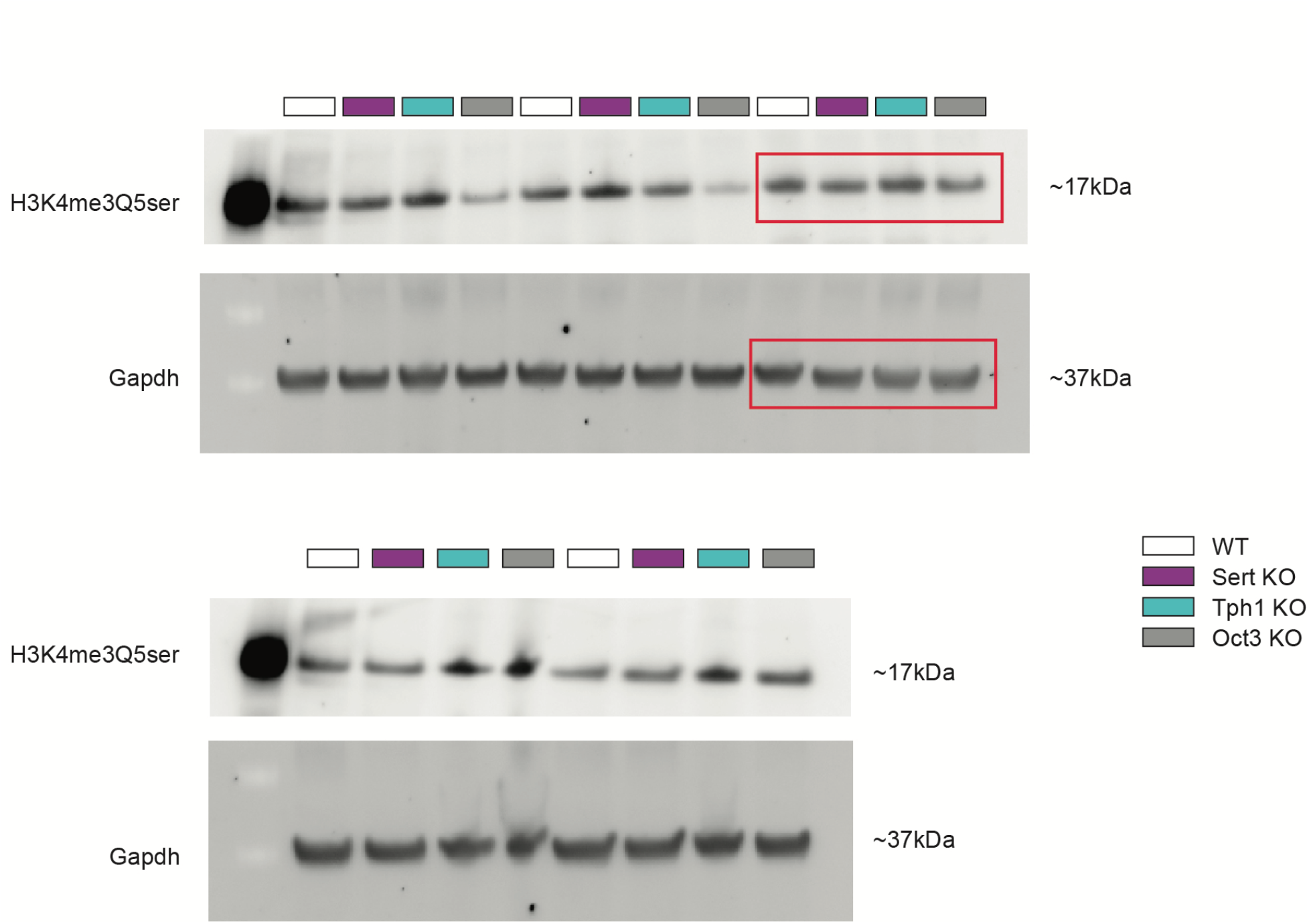
Western blots used for the quantification of H3K4me3Q5ser in E12.5 brain tissues of WT, Tph1 KO, Sert KO, and Oct3 KO in. Figure 4D. Red rectangles indicate the representative blots displayed in the main figure.

## Supplementary Table Captions

Excel file including Supplementary Tables 1-28, which contain analyses of ChIP-seq and RNA-seq from placental and brain tissues.

**Supplementary Table 1:** Developmental placenta H3K4me3Q5ser ChIP-seq, DiffBind results (E9.5 male vs E17.5 male): Fig. 1E-J, Supplementary Fig. 3A

**Supplementary Table 2:** Developmental placenta H3K4me3Q5ser ChIP-seq, DiffBind results (E9.5 female vs E17.5 female): Fig. 1E, H, I, Supplementary Fig. 3B-C

**Supplementary Table 3:** Developmental placenta H3K4me3Q5ser ChIP-seq, DiffBind results (E9.5 male vs E9.5 female): Fig. 1E, Supplementary Fig. 2A

**Supplementary Table 4:** Developmental placenta H3K4me3Q5ser ChIP-seq, DiffBind results (E12.5 male vs E12.5 female): Fig. 1E, Supplementary Fig. 2B

**Supplementary Table 5:** Developmental placenta H3K4me3Q5ser ChIP-seq, DiffBind results (E17.5 male vs E17.5 female): Fig. 1E, Supplementary Fig. 2C

**Supplementary Table 6**: Developmental placenta bulk RNA-seq, normalized counts table

**Supplementary Table 7**: Developmental placenta bulk RNA-seq, DESeq2 results (E9.5 male vs E17.5 male): Fig. 1I, Supplementary Fig. 3D-E

**Supplementary Table 8**: Developmental placenta bulk RNA-seq, DESeq2 results (E9.5 female vs E17.5 female): Fig. 1I, Supplementary Fig. 3D-E

**Supplementary Table 9**: Developmental placenta functional annotation analysis, Gene Ontology Biological Processes (E9.5 male vs E17.5 male): Fig. 1K

**Supplementary Table 10**: Developmental placenta functional annotation analysis, Reactome (E9.5 female vs E17.5 female): Fig. 1K

**Supplementary Table 11:** Transgenic placenta H3K4me3Q5ser ChIP-seq, DiffBind results (WT vs Sert KO): Fig. 3A-C, Supplementary Fig. 7C-D

**Supplementary Table 12:** Transgenic placenta H3K4me3 ChIP-seq, DiffBind results (WT vs Sert KO): Fig. 3A, 3C

**Supplementary Table 13:** Transgenic placenta H3K4me3Q5ser ChIP-seq, DiffBind results (WT vs Tph1 KO): Fig. 3A, 3C, Supplementary Fig. 7A

**Supplementary Table 14:** Transgenic placenta H3K4me3 ChIP-seq, DiffBind results (WT vs Tph1 KO): Fig. 3A, 3C, Supplementary Fig. 7A

**Supplementary Table 15:** Transgenic placenta H3K4me3Q5ser ChIP-seq, DiffBind results (WT vs Oct3 KO): Fig. 3A, 3C, Supplementary Fig. 7B

**Supplementary Table 16:** Transgenic placenta H3K4me3 ChIP-seq, DiffBind results (WT vs Oct3 KO): Fig. 3A, 3C, Supplementary Fig. 7B

**Supplementary Table 17**: Transgenic placenta functional annotation analysis, Gene Ontology Biological Processes (WT vs Sert KO): Fig. 3G

**Supplementary Table 18**: Transgenic placenta functional annotation analysis, Gene Ontology Biological Processes (WT vs Tph1 KO): Fig. 3G

**Supplementary Table 19**: Transgenic placenta functional annotation analysis, Gene Ontology Biological Processes (WT vs Oct3 KO): Fig. 3G

**Supplementary Table 20**: Transgenic brain bulk RNA-seq, normalized counts table

**Supplementary Table 21**: Transgenic brain bulk RNA-seq, DESeq2 results (WT vs Sert KO): Fig. 4E-F

**Supplementary Table 22**: Transgenic brain bulk RNA-seq, DESeq2 results (WT vs Tph1 KO): Fig. 4E-F

**Supplementary Table 23**: Transgenic brain bulk RNA-seq, DESeq2 results (WT vs Oct3 KO): Fig. 4E-F

**Supplementary Table 24**: Transgenic brain bulk RNA-seq functional annotation analysis, Gene Ontology Biological Processes (WT vs Sert KO): Fig. 4G

**Supplementary Table 25**: Transgenic brain bulk RNA-seq functional annotation analysis, Reactome (WT vs Sert KO): Fig. 4G

**Supplementary Table 26**: Transgenic brain bulk RNA-seq functional annotation analysis, Gene Ontology Biological Processes (WT vs Oct3 KO): Fig. 4G

**Supplementary Table 27**: Transgenic brain bulk RNA-seq functional annotation analysis, Reactome (WT vs Oct3 KO): Fig. 4G

**Supplementary Table 28**: Transgenic brain bulk RNA-seq functional annotation analysis, Reactome (WT vs Tph1 KO): Fig. 4G

## Acknowledgements

This work was partially supported by grants from the National Institutes of Health: R01 MH116900 (I.M.), F32 MH126534 (J.C.C.), F31 NS132558 (A.M.C.), as well as funds from the Howard Hughes Medical Institute (I.M.). NA and MB were supported by the EU H2020 MSCA ITN projects “Serotonin and Beyond” (N 953327). All schematics were created with Biorender.com.

## Author Contributions

J.C.C. and I.M. conceptualized the study. J.C.C. performed the experiments, collected and analyzed the data. N.A. and M.B. provided mouse tissues. J.C.C., A.M.C., A.R. and L.S. performed the bioinformatics analyses. J.C.C. and I.M. wrote the manuscript.

